# Microglia become hypofunctional and release metalloproteases and tau seeds after phagocytosing live neurons with P301S tau aggregates

**DOI:** 10.1101/2021.02.26.433088

**Authors:** Jack H. Brelstaff, Matthew Mason, Taxiarchis Katsinelos, William A. McEwan, Bernardino Ghetti, Aviva M. Tolkovsky, Maria Grazia Spillantini

## Abstract

The microtubule-associated protein tau aggregates in multiple neurodegenerative diseases, causing inflammation and changing the inflammatory signature of microglia by unknown mechanisms. We have shown that microglia phagocytose live neurons containing tau aggregates cultured from P301S tau transgenic mice due to neuronal tau aggregate-induced exposure of the ‘eat me’ signal phosphatidylserine. Here we show that after phagocytosis, microglia become hypophagocytic while releasing seed-competent insoluble tau aggregates. These microglia activate acidic β-galactosidase, and release senescence-associated cytokines and matrix remodeling enzymes alongside tau, indicating a senescent phenotype. In particular, the marked NFκB-induced activation of matrix metalloprotease 3 (MMP3/stromelysin1) was replicated in the brains of P301S mutant tau transgenic mice, and in human brains from tauopathy patients. These data show that microglia that have been activated to ingest live neurons with tau aggregates behave hormetically, becoming hypofunctional while acting as vectors of tau aggregate spreading.

## Introduction

Aggregation of the protein tau from a soluble unfolded state to an insoluble β-sheet enriched filamentous structure underlies numerous human neurodegenerative diseases known as tauopathies (*1*). These include Alzheimer’s disease, various frontotemporal dementias, Pick’s disease, progressive supranuclear palsy, corticobasal degeneration, chronic traumatic encephalopathy, and argyrophilic grain disease. The presence of intraneuronal aggregates of tau best correlate with the neuronal cell death that is associated with the clinical signs and symptoms of disease (*2*). The mechanism of cell death in tauopathy remains unclear, however several studies implicate microglia in non-cell-autonomous routes (*3–8*).

Microglia are the resident immune cells of the brain and in neurodegeneration become pathologically activated leading to proliferation and release of cytotoxic cytokines. Microglia have high phagocytic potential and remove synapses during developmental pruning (*9–1*) via phosphatidylserine (PtdSer) (*12, 13*) as well as apoptotic bodies (*14*). Recent evidence shows that microglia are capable of aberrantly phagocytosing synapses and viable neurons in the adult mouse brain (*15–17*) and this may be increased in Alzheimer’s disease and frontotemporal dementia (*18, 19*). We have previously shown that inflammation is present in human mutant P301S tau mice (P301S tau) (*20*) and that live cultured dorsal root ganglion neurons (DRGn) from P301S tau mice display phosphatidylserine on the external leaflet of their plasma membranes which recruits cocultured microglia that phagocytose them while still viable (*17*). This process is accompanied by secretion of the opsonin milk fat globule EGF 8 (MFGE8) and nitric oxide production, and leads to the transfer of tau aggregates inside the phagocytosed neurons to the microglia (*17*). Microglia have also been shown to phagocytose extracellular tau and become activated to phagocytose neurons in a PKC dependant manner (*4*). These data implicate microglia in the pathological loss of neurons and synapses in tauopathy by intensifying phagocytic activity to damaging levels.

Tau misfolds into distinct fibrillar forms depending on the specific tauopathy (*21*). Different filamentous forms of tau can template specific pathological conformations onto naïve monomers through a mechanism of prion-like spreading (*22*). Analysis of Braak stage progression suggests that misfolded tau is released as seeds through synaptic connections because anatomically connected regions progressively develop pathology (*23, 24*). Microglia have also been implicated in the spreading of tau pathology (*25*) by releasing exosomes containing previously phagocytosed tau that is capable of propagating *in vivo* (*26*). A 40% microglial elimination by CSF1R inhibition reduced neurodegeneration in the spinal cords of human P301S mutant transgenic mice (*27*) but a similar 30% reduction in microglial number did not affect the degree of tau pathology in the cortex of a human P301L tau mutant transgenic mouse where the transgene is expressed under the Cam Kinase II promoter (*28*). If microglia release tau seeds after phagocytosing either a tau aggregate-containing neuron or extracellular tau released at a synapse, they could act as vectors of tau spreading in the local environment.

Here we report that *in vitro* microglia that have phagocytosed living neurons with insoluble filamentous tau aggregates contain tau foci, and release insoluble tau aggregates into the conditioned medium. Tau released from such microglia seeds new tau aggregates that are hyperphosphorylated in a reporter system consisting of HEK-P301S-venus expressing cells. Microglia cocultured with tau aggregate-containing neurons are hypophagocytic towards both latex beads and a second exposure to tau aggregate-containing neurons that expose the phagocytic signal phosphatidylserine. These microglia, recultured following coculture, also show an increase in β-galactosidase (SA β-gal) activity, a marker of cellular senescence. Furthermore, microglia cocultured with aggregated P301S tau-containing neurons secrete a unique signature of proteins that differs from that of LPS-stimulated microglia. Among these secreted proteins, the active form of matrix metalloprotease 3 (MMP3) is not only highly induced in microglia by coculture with tau aggregate-containing neurons, but is also increased in the brains of P301S tau transgenic mice and in brains of patients with tauopathies. Hence, microglia that have phagocytosed neurons containing tau aggregates enter a unique hypofunctional state that resembles aspects of senescence.

## Results

### Microglia that have phagocytosed P301S tau neurons containing tau aggregates release tau

DRGn cultures prepared from 5-month-old P301S tau (5M P301S) mice (*29, 30*) were cocultured for 4 days with wild type microglia obtained from C57Bl6J mice (C57) to induce phagocytosis of live neurons containing filamentous tau aggregates as described previously (*17*). Neuronal tau aggregates that were prelabelled with pFTAA before addition of microglia (*31*) were visible as foci of 3-5 μm diameter and smaller puncta (Figure 1A). Conditioned media (CM) were assayed for tau by ELISA after 4 days from either monocultured neurons or microglia, or from cocultured microglia with neurons, or from microglia re-isolated after coculture and cultured alone for a further 4 days (Figure 1B). No measurable tau was detected in CM from monocultured P301S DRGn containing tau aggregates, C57 wild type DRGn or naïve monocultured microglia. However, 4.54 ± 0.49 pg/ml tau was detected in the CM from P301S DRGn-microglia cocultures (average ± SD, p=0.012, one sample *t* test), and continued to be present even when microglia were re-isolated from the cocultures and cultured alone for a further 4 days (8.4 ± 1.3 pg/ml, p=0.185 compared to P301S DRGn-microglia cocultures unpaired *t* test), suggesting that the source of tau was specifically microglia that had phagocytosed neurons with tau aggregates (Figure 1B). No significant differences in LDH release were detected in the CM across all conditions, demonstrating that tau was not released due cell death and lysis (p=0.069, one way ANOVA) (Figure 1B).

**Fig. 1.**
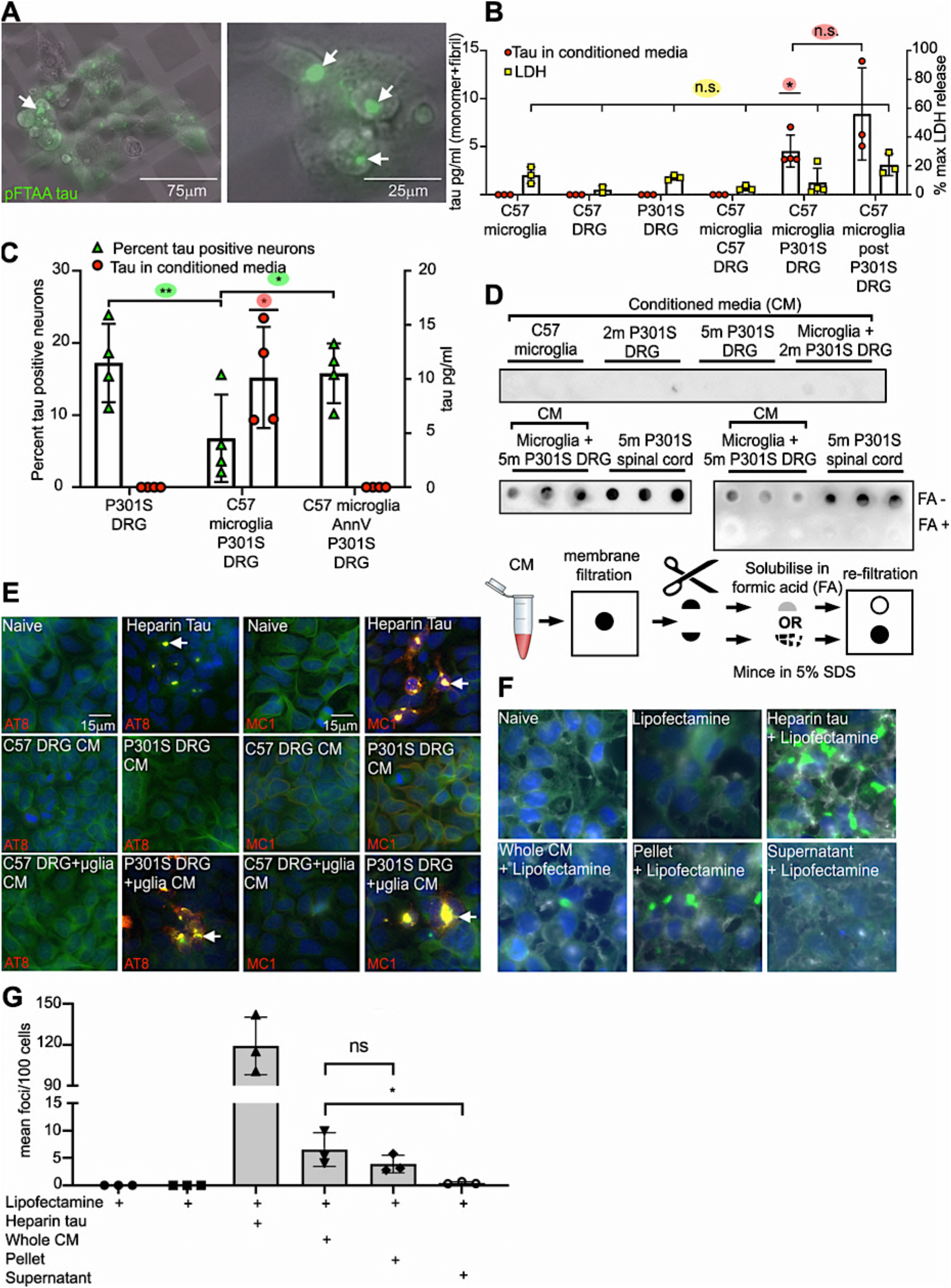
Microglia that have phagocytosed P301S DRGn containing tau aggregates release insoluble tau seeds into the CM. **A**) 5M P301S DRGn with tau aggregates prelabelled with pFTAA are transferred to microglia after phagocytosis. Microglia were purified after 4 days of coculture, replated and live imaged. Images show combined phase and fluorescence. **B**) Tau is only released into the CM under condition of coculture with 5M P301S DRGn (p=0.012, one sample t test). Human-specific tau detection is by ELISA. Tau continues to be released into the CM from re-isolated microglia post coculture with 5M P301S DRGn (p=0.185 vs coculture, unpaired t test). No tau is detected in CM from monocultures of 5M P301S DRGn, 5M C57 DRGn, or C57 microglia, or cocultures of 5M C57 DRGn and microglia. Lactate dehydrogenase (LDH) activity is not significantly different in CM across all conditions, indicating that released tau is not due to lysed cells (p=0.069, one way ANOVA). N= 3 independent biological replicates ± SD. **C**) Release of tau requires phagocytosis by microglia. Number of tau-positive 5M P301S DRGn (% HT7/βIII tubulin) is reduced after coculture with microglia (p=0.0048, repeated measures ANOVA), commensurate with an increase in tau in the CM (p=0.0225, one sample t test), whereas blocking phagocytosis by masking PtdSer with AnnV prevents neuronal loss (n.s. vs monocultured P301S DRGn, p=0.0309 vs coculture but no AnnV, repeated measures ANOVA), and prevents the release of tau into the CM (n.s. vs monocultured P301S DRGn). N= 4 independent biological replicates ± SD. **D**) Microglia cocultured with 5M P301S DRGn release insoluble tau aggregates into the CM. Tau in the CM is captured on a filter trap. Top panel: monocultures of DRGn (5M C57, 2M P301S, and 5M P301S), and coculture of C57 microglia with 2M P301S which do not contain filamentous tau aggregates; Middle left panel, coculture of microglia with 5M P301S DRGn, and spinal cord extracts from 5M P301S as positive controls. Note that these panels are segmented from the same membrane and developed simultaneously (Supplementary Figure 1). Middle right pane: Refiltration of captured tau solubilised in formic acid (+FA) leads to loss of signal but minced without FA retains signal. The procedure is schematically represented in the bottom panel. **E**) Tau in CM seeds new aggregates in the reporter system HEK-P301S-venus cells. The tauseeded aggregates contain hyperphosphorylated and conformationally altered tau as shown by AT8 and MC1 antibody staining. **F & G**) Aggregated P301S-venus foci are obtained with the 100,000x*g* pellet but not the supernatant of CM after ultracentrifugation (p=0.0429, Friedman test). Representative images and quantification of fractionated CM seeded aggregation in HEK-P301S-venus cells. Lipofectamine was added to increase the efficiency of the uptake. N=3 independent biological replicates ± SD. Significant differences are shown as * p<0.05, ** p<0.01.

To further confirm that microglia were the source of tau in the CM post-phagocytosis, PtdSer in neurons was masked with Annexin V (AnnV) before coculture (*17*). In the absence of AnnV, the proportion of P301S DRGn neurons (% HT7 positive/βIII tubulin positive neurons) was significantly reduced ~2-fold when cocultured with microglia (p=0.0048, repeated measures ANOVA), but this loss was prevented by addition of AnnV (p=0.0309 vs without AnnV, p=0.378 vs naïve) (Figure 1C). In keeping with a requirement for engulfment of P301S DRGn for tau release by microglia, 10 ± 4.8 pg/ml tau was detected in the CM of cocultured P301S DRGn and microglia but no tau was detected in the CM from cocultures when phagocytosis and loss of tau positive neurons was prevented by adding AnnV (p=0.0225 comparing ± AnnV, one sample *t* test) (Figure 1C). Thus, microglia are the source of tau released into the CM.

### Tau in the CM is insoluble and seeds tau aggregation

To investigate whether released tau is monomeric, or remains as larger order aggregates similar to those found in the neurons from 5M P301S mice (*29, 31*), oligomers in the CM were solubilised with SDS (5%) and the preparation was filtered through a 0.2 μm cellulose acetate membrane (pre-soaked in 5% SDS) under vacuum, a method which traps insoluble tau aggregates from AD brains (*32*). Sarkosyl-insoluble tau aggregates from the spinal cord of 5M P301S mice (*17, 30*) were used as a positive control. CM from cocultured 5M P301S DRGn and microglia contained tau species large enough to be captured on the membrane (Figure 1D) but no tau was detected in CM from monocultures of microglia, 5M P301S DRGn, or 2M P301S DRGn (which do not contain aggregated tau (*29*)), or from cocultured 2M P301S DRGn with microglia. To further demonstrate that the tau coming from the coculture CM and trapped on the filter was insoluble, each individual filtrate was cut in half as illustrated in Figure 1D; one half was treated with 70% formic acid (FA), which solubilises tau aggregates (*31*), dialysed to remove FA, and re-filtered alongside extracts from the other untreated half of the filter, which was only minced and soaked in 5% SDS. FA treated samples showed no signal whereas the extracts from the minced samples were still trapped on the membrane (Figure 1D), indicating that the tau assemblies trapped on the membrane are insoluble aggregates.

To assay whether tau released from microglia into the CM has aggregation-inducing properties, we used the HEK-P301S-venus cell reporter system (*33*). HEK-P301S-venus cells were incubated with CM for 24 h, fixed and immunostained with AT8 to detect hyperphosphorylated tau, or with MC1, which is a conformational antibody that solely detects misfolded tau (*17, 34, 35*). Addition of heparin-tau aggregates was used as a positive control. CM from cocultured 5M P301S DRGn and microglia seeded AT8- and MC1-positive aggregates (Figure 1E) whereas CM from monocultures of C57 or P301S DRGn or C57 DRGn cocultured with microglia showed no seeding capability. To further determine the characteristics of the tau seeds released into CM, the CM was ultracentrifuged at 100,000x*g* for 30 minutes to separate soluble oligomeric species from insoluble aggregates. The pellet was resuspended to the original volume of the CM, and both fractions were added to HEK-P301S-venus cells for 4 days in the presence of lipofectamine to enhance seed uptake (*33*). The pellet fraction retained a similar aggregation activity to that of whole CM (p=0.662, Friedman test) whereas the supernatant produced significantly fewer foci (p=0.0429 Friedman test) (Figure 1F & G). Thus, microglia that have phagocytosed neurons with tau aggregates release tau as species that are competent to seed insoluble tau aggregates in recipient cells.

### Re-isolated microglia from P301S tau neuron cocultures are hypophagocytic

We next investigated whether and how phagocytosis of neurons with tau aggregates affects microglial function. To test their phagocytic capacity, microglia were isolated from cocultures with 5M P301S DRGn and were cocultured again with a new monoculture of 5M P301S DRGn. While coculture with fresh C57 microglia caused a significant loss of tau neurons (p=0.0013, one way ANOVA), no significant loss of tau aggregate-containing neurons was detectable in coculture of re-isolated C57 microglia (p=0.9903, one way ANOVA) (Figure 2A), thus indicating that following ingestion of DRGn containing tau aggregates the microglia become hypophagocytic and unable to ingest other DRGn with tau aggregates.

**Fig. 2.**
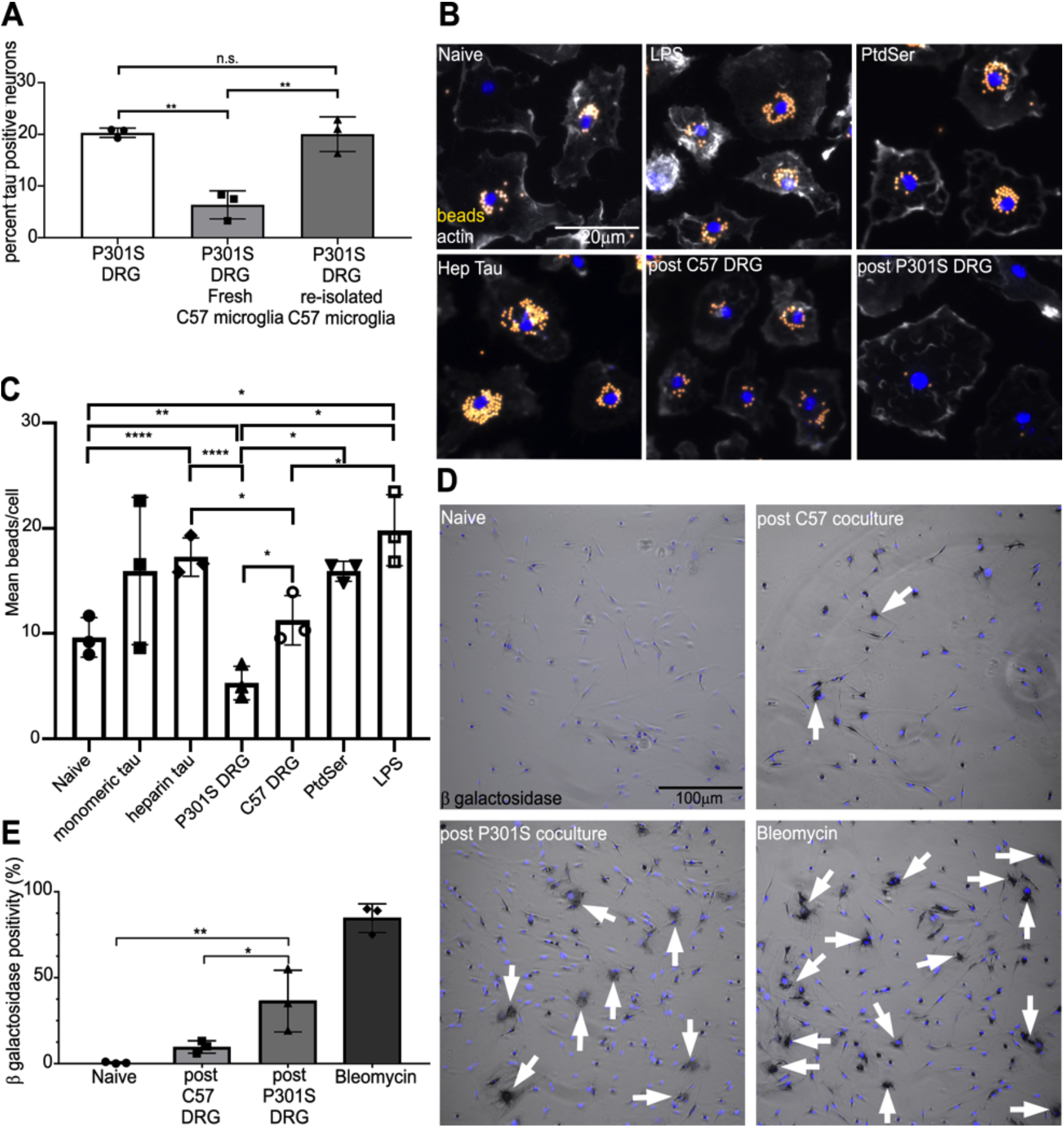
Re-isolated microglia from cocultures of 5M P301S DRGn and microglia are hypophagocytic and display increased SA-β-gal activity. **A**) C57 microglia re-isolated after previous coculture with 5M P301S DRGn fail to remove tau aggregate-containing neurons compared to fresh C57 microglia (5M P301S DRGn alone vs neurons with fresh microglia p=0.0013, or vs re-isolated microglia, p=0.9903, one way ANOVA). N= 3 independent biological replicates ± SD. **B & C**) Re-isolated microglia from coculture with 5M P301S DRGn show a significant reduction of latex bead uptake compared to naïve microglia in monoculture (p=0.007) or microglia re-isolated from coculture with 5M C57 DRGn (p=0.0239, repeated measures one way ANOVA); treatment of naïve microglia with PtdSer liposomes does not reduce bead uptake (n.s.), while treatment with heparin-assembled tau significantly stimulates bead uptake (p<0.0001). N=3 independent biological replicates ± SD. Significant differences are shown as * p<0.05, ** p<0.01, *** p<0.001, **** p<0.0001. **D & E**) The senescence marker SA-β-gal is significantly increased in re-isolated microglia from cocultures with 5M P301S DRGn compared to cocultures with 5M C57 DRGn (p=0.0168) or naïve microglia (p=0.0025, one-way ANOVA and Sidak post hoc). The senescence inducer Bleomycin is used as a positive control. N= 3 independent biological replicates ± SD. Significant differences are shown as * p<0.05, ** p<0.01.

Phagocytosis of 5M P301S neurons is dependent on the specific exposure of the phagocytic signal phosphatidylserine (PtdSer) on their membranes (*17*). To investigate whether inhibition of microglial phagocytosis was independent of exposure to living P301S neurons and a generalised effect, re-isolated microglia were incubated with carboxylate-modified latex beads. Similarly to the suppressed phagocytosis after exposure to 5M P301S tau DRGn, re-isolated microglia from 5M P301S tau DRGn cocultures phagocytosed significantly fewer beads than microglia reisolated from C57 DRGn cocultures (p=0.0239, one way ANOVA with repeated measures) or naïve microglia (p=0.007), which had equivalent bead uptake to the C57 DRGn cocultures (p=0.1507) (Figure 2B, quantified in C). In contrast to the suppression of phagocytoses induced in microglia by the coculture with 5M P301S DRGn, exposure to heparinised-tau increased bead phagocytosis significantly (p=<0.0001), suggesting that it was the ingestion of the neurons containing tau aggregates, and not the exposure to released aggregated tau, that causes this hypofunctionality. Monomeric tau also appeared to increase average bead uptake, but the increase was not significant (p=0.5487). Exposure to PtdSer liposomes, to mimic the PtdSer exposure on 5M P301S DRGn, caused a similar bead uptake to naïve microglia (p=0.557), which was significantly higher compared to the uptake by microglia that had been re-isolated from the cocultures with 5M P301S DRGn (p=0.0139). These results indicate that the hypophagocytosis by microglia post P301S DRGn coculture was not related to PtdSer signalling. LPS, which was added as a positive control, increased bead uptake by the microglia by about 2-fold compared to the uptake by naïve microglia (p=0.028), similar to the uptake induced by monomeric and heparin tau. Thus, microglia that have phagocytosed P301S DRGn with tau aggregates become hypophagocytic independently of PtdSer exposure.

### Hypophagocytic microglia activate senescence-associated acidic β galactosidase

Since we had observed a loss of microglial function due to coculture with tau aggregate containing neurons, and pathological tau is associated with deleterious microglial cellular senescence in a mouse model of tauopathy and in AD (*36, 37*), we investigated whether senescence was induced in these hypofunctional microglia. We first tested whether microglia reisolated from coculture with 5M P301S DRGn activate the senescence marker acidic β galactosidase (SA-β gal). There were significantly more β galactosidase-positive microglia in cultures re-isolated from 5M P301S DRGn cocultures compared to microglia re-isolated from cocultures with C57 DRGn (~3-fold, p=0.0168, one way ANOVA) or naïve microglia (p=0.0025) (Figure 2D, quantified in E). As a positive control for the assay, we exposed naïve microglia to bleomycin, which is a well-established inducer of cellular senescence (*38, 39*). Bleomycin (50 μg/ml, 12 hours) upregulated acidic β galactosidase activity about 8-fold compared to microglia cocultured with C57 DRGn. Thus, microglia that phagocytosed 5M P301S DRGn with tau aggregates are not only hypophagocytic, but also appear to be have acquired a senescence-like state.

### Hypophagocytic microglia release a specific senescence-like protein profile highly enriched in MMP3

Microglia cocultured with 5M P301S DRGn release MFGE8 and nitric oxide into the CM, signals essential for microglial phagocytosis of these neurons (*17*). To investigate whether microglia that have phagocytosed P301S DRGn produce signals that may account for their hypofunctional post-phagocytic state, we screened the CM for the expression of 86 mouse proteins (consisting of cytokines, chemokines, growth factors, and matrix-modifying proteases) across multiple treatment conditions using the membrane Proteome Profiler™ Cytokine array. Supplementary Figure 2 shows representative membranes from each culture condition while Supplementary Figure 2 B shows quantification of relative intensities normalised to internal standards across the entire membrane. DRGn monocultures from 5M C57 mice or 5M P301S mice produced no detectable cytokines but naïve microglial monocultures produced a basal expression pattern that includes CCL2, CCL6, Chemerin, IGFBP-6, m-CSF1, Osteopontin, Serpin E1, and VEGF. LPS treatment of naïve microglia, used to profile an inflammatory phenotype, induced a classical inflammation-associated profile including strong TNFα expression. In contrast, no pro-inflammatory cytokines like TNFα were detected in CM from 5M P301S neuron-microglial cocultures. Instead, a different profile of proteins was released.

Submitting the full array of proteins to STRING analysis (*40*) identified MMP3 plus 14 putative partners (MMP3, MMP9, CXCL2/MIP2, CXCL1/KC, MMP2, VEGF, Endostatin, IGFBP3, Pentraxin2, CXCL10, CCL5/Rantes, CXCL5, CXCL9, CXCL13, CCL20). Of these, CXCL5, CXCL9, CXCL13, and CCL20 showed no expression across any of the conditions assayed, and were therefore excluded from further analysis. The relative expression values and row Z scores for the remaining 11 proteins are shown in Figure 3A, and their functional interaction network generated by the STRING analysis shown in Figure 3B. Principal component analysis (PCA) of the 11 proteins showed that the subset of proteins released by microglia cocultured with 5M P301S DRGn formed a unique group, distinct from LPS-treated microglia, or those cocultured with C57 DRGn, or 2M P301S DRGn (Figure 3 C), as also corroborated by the biweight midcorrelation analysis (Figure 3 D). A loading plot (Figure 3 E) shows that the metalloproteases were major contributors to PCA1.

**Fig. 3.**
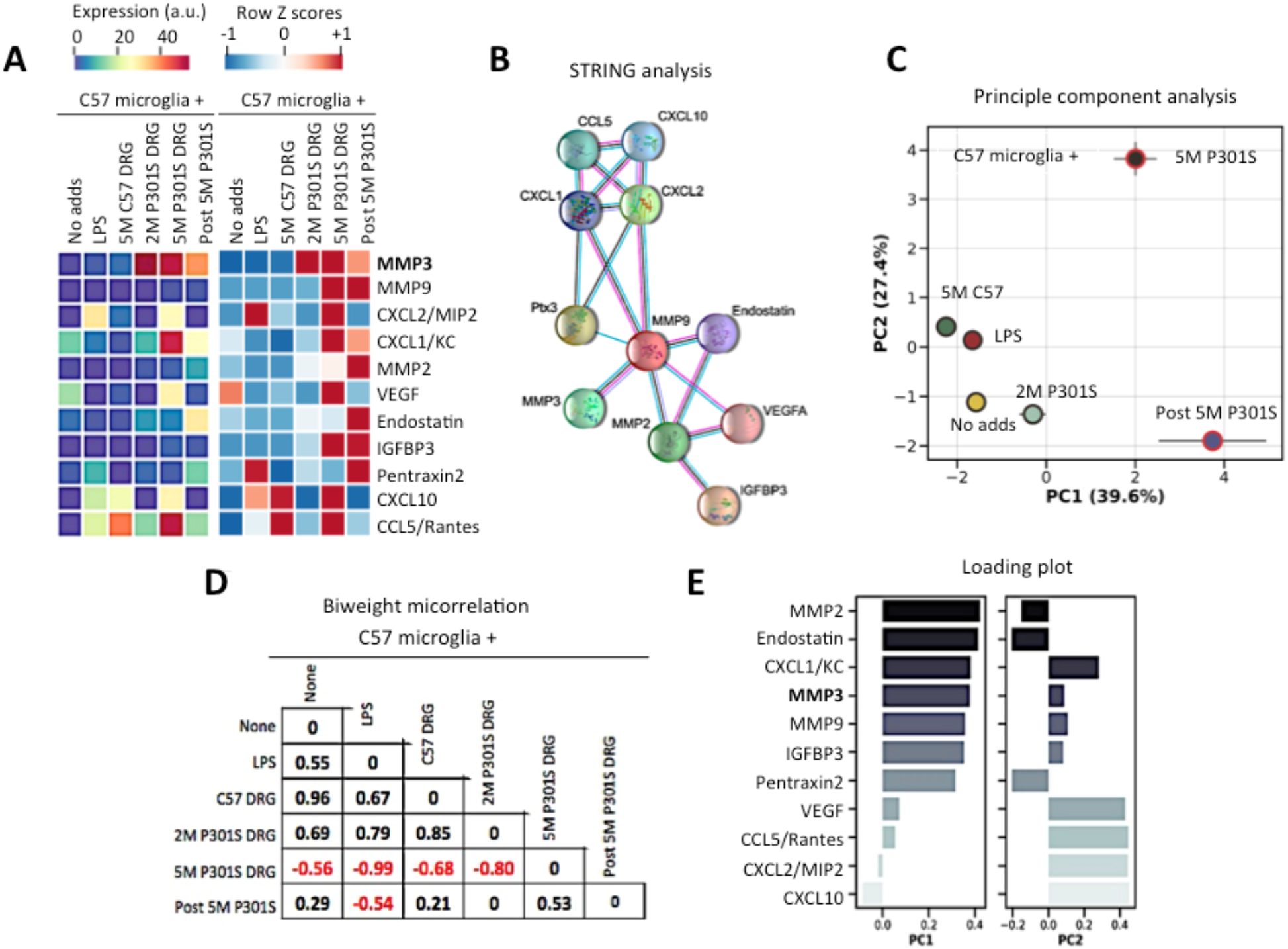
Microglia cocultured with 5M P301S DRGn secrete a unique protein profile. **A)** Comparative heat-map analysis of protein expression of a family of proteins that are highly enriched in CM from cocultures of microglia and 5m P301S DRGn or microglia re-isolated from these cultures, but not in naïve microglia or those cocultured 5M C57 DRGn or 2M P301S DRGn (with no filamentous tau aggregates). **B**) Functional interaction network between the 11 proteins identified by STRING analysis. **C)** PCA plot of proteins presented in panel A (mean ± SD, n=3 independent biological replicates). **D**) Biweight midcorrelation matrix of the PCA data showing a strong negative correlation between the condition of coculture of microglia with 5M P301S DRGn and all other experimental conditions except for the post-cultured microglia. **E**) Loadings plot of the two principal components, showing that MMPs dominate the PC1 axis whereas PC2 is dominated by the cytokines CXCL1, −2, −10, and CCL5.

### MMP3 is upregulated in the CM of cocultured 5M P301S DRGn and microglia and brains of P301S mice and human tauopathies

Because MMP3 was the most prominently-upregulated protein (Figure 3 A, Supplementary Figure 2A), we examined whether its expression is also increased in mouse and human brains associated with tauopathies. Study of MMP3 expression in the frontal cortex of 5M C57, 2M P301S, and 5M P301S mice by immunoblotting showed two bands corresponding to the inactive zymogen (pro form) and active (mature) MMP3 (*41, 42*). Both the pro and active forms were present in wild type mouse brain but there was a highly significant conversion from the inactive to the active form in 5M P301S mice (p<0.001, one-way ANOVA) (Figure 4 A & B). There was an increase in active MMP3 in 2M P301S mouse brain as well, but this did not reach significance when measured by semi-quantitative densitometry (p=0.9120) (Figure 4 B). To investigate whether this upregulation of MMP3 was relevant to human disease, tissue lysates from the frontal cortex of non-demented controls and patients with FTDP-17T with +3 or P301L MAPT mutations, or Pick’s disease, and from the midbrain of Progressive Supranuclear Palsy (PSP) patients were immunoblotted for MMP3 (Figure 4 C & D). FTD with TDP-43 pathology due to a C9orf72 expansion mutation was included to investigate whether MMP3 upregulation is specific to pure tauopathies. The amount of active MMP3 was barely detected in extracts from control brains, but there was a substantial increase in the active/mature form in all the samples from neurodegenerative diseases: FTDP-17T with the +3 and the P301L mutations were elevated 4.7-fold (p=0.013) and 4.5-fold (p=0.043), respectively (ANOVA p=0.04, 2-tailed t test compared to control) while the ratios were slightly decreased for C9orf72 (1.78-fold, p=0.003), PSP (2.45-fold, p=0.019), and Pick’s disease (3.07-fold, p=0.025) (ANOVA p=0.012, 2-tailed t test compared to control) (Figure 4 D). Hence, active MMP3 is a candidate marker of advanced tauopathies and at least one related dementia.

**Fig. 4.**
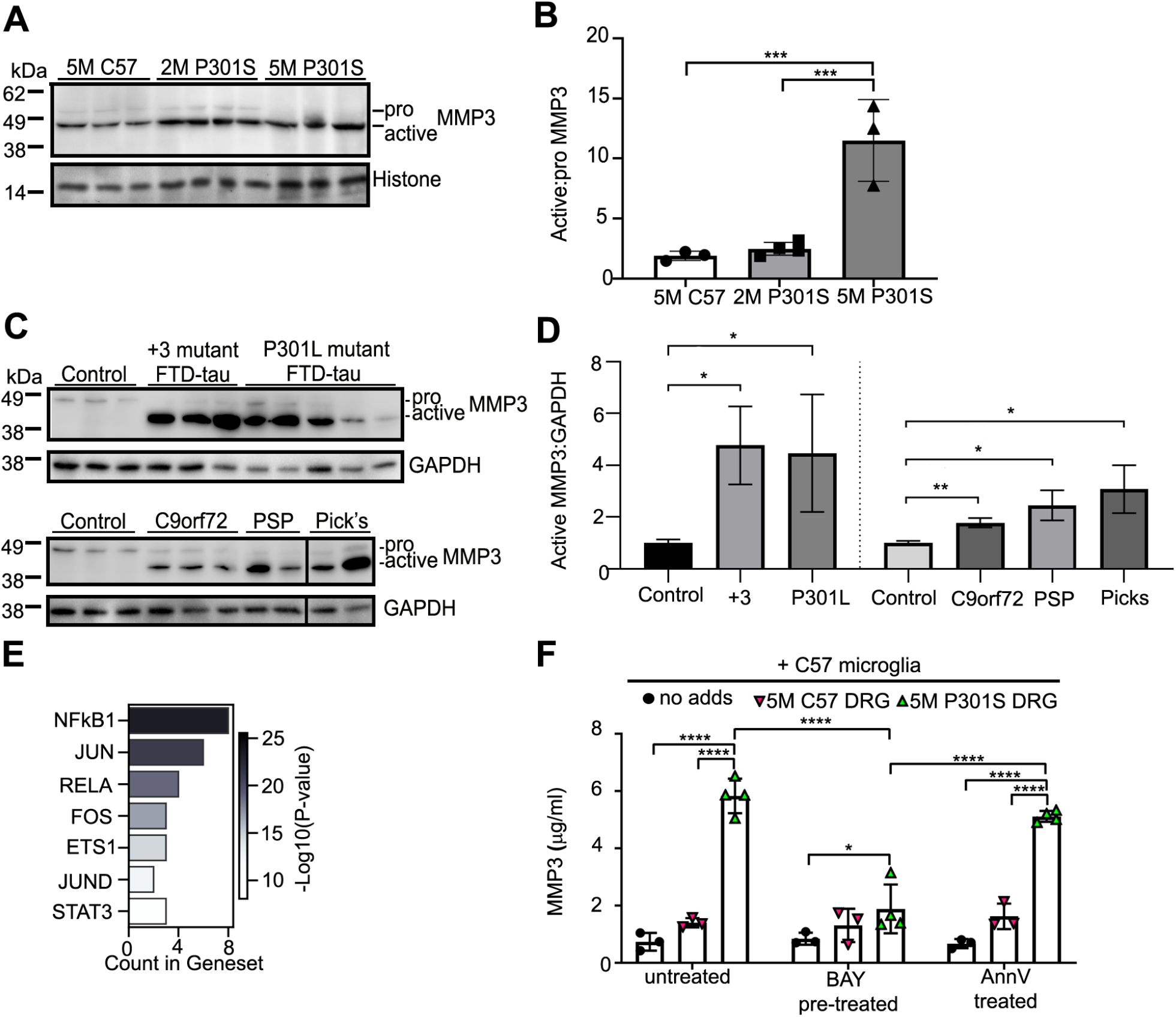
MMP3 is converted to its active form in transgenic P301S mice and human neurodegenerative disease and is under transcriptional control of NFκB. **A**) Immunoblot of pro- and active MMP3 expression in the frontal cortex of C57 wild type and P301S transgenic mouse brain. **B**) Ratio of active/pro forms of MMP3 showing significantly more active MMP3 is expressed in the brains of 5M P301S mice. **C**) MMP3 expression in the frontal cortex of controls, +3 and P301L mutant FTDP-17T cases, FTD with C9orf72 mutation cases, Pick’s disease, and midbrain from PSP cases. **D**) Ratio of active MMP3/GAPDH intensities showing greater active MMP3 expression in neurodegenerative disease vs healthy control. **E**) The number of proteins identified in the transcriptional regulation network using the Enrichr programme; the shaded bar denotes −log10 transformation of adjusted P-values that were computed by Fisher’s exact test within Enrichr. **F**) Pre-treatment of microglia with the NFκB pathway inhibitor BAY11-8072 significantly prevents the expression of MMP3 in cocultures of P301S aggregate-containing neurons and microglia. AnnV masking of Ptdser does not prevent MMP3 production. *p<0.05; *p<0.01,; ***p<0.001;****p<0.0001.

### NFkB regulates MMP3 upregulation

Previous studies have implicated the NFκB signalling pathway in the regulation of several of the proteins identified in the CM from 5M P301S DRGn-microglia cocultures (*43*). Submission of the 11 proteins identified in Figure 3A to the open-source gene set enrichment analysis database Enrichr (*44*) showed that 8 proteins, including MMP3, were predicted to be under the transcriptional control of NFκB1, and 4 proteins were predicted to be under the transcriptional control of RelA (NFκB p65 subunit) (Figure 4 E). We therefore investigated whether NFκB controls the production of MMP3 in our experimental paradigm. Microglia were preincubated with the NFκB pathway inhibitor BAY11-8072 (*45*) for 1 h, after which they were cocultured with 5M C57 or 5M P301S tau DRGn. CM was collected after 24 h and MMP3 was measured by ELISA. BAY11-8072 significantly reduced the amount of MMP3 in the CM produced by coculture of 5M P301S DRGn and microglia (p<0.0001, two-way ANOVA Tukey post hoc) (Figure 4 F). BAY11-8072 induced no change in the amount of MMP3 detected in co culture with 5M C57 DRGn. To investigate whether MMP3 expression induced by 5M P301S tau DRGn in microglia was due to phagocytosis of the neurons, phagocytosis was prevented by preincubating the neurons with annexin V (AnnV). Unlike BAY11-8072, AnnV blockade of phagocytosis did not reduce the production of MMP3 (p<0.0001), indicating that phagocytosis-related hypofunctionality and NFκB-dependent MMP3 production are activated by different effectors emanating from the 5M P30S-DRGn. These results show that while hypofunctionality is a consequence of the physical act of phagocytosis linked to PtdSer, MMP3 activation appears to depend on additional signals that are PtdSer-independent.

## Discussion

Inflammation and microgliosis are a common feature of many neurodegenerative diseases and microglial phagocytosis is suggested to be highly important in disease pathogenesis with the discovery of phagocytosis genes as genetic risk factors (*46*). Here we report that after phagocytosis of neurons with hyperphosphorylated and filamentous P301S tau aggregates, microglia undergo profound changes: they release insoluble forms of tau that can seed new tau aggregates, they become hypophagocytic, and they display several elements of senescence, including senescence-associated acidic β galactosidase activity and release of senescence-associated proteins, some of which are under NFκB transcriptional control. A scheme illustrating how these events might lead to a vicious cycle of neurodegeneration is shown in Figure 5.

**Fig. 5.**
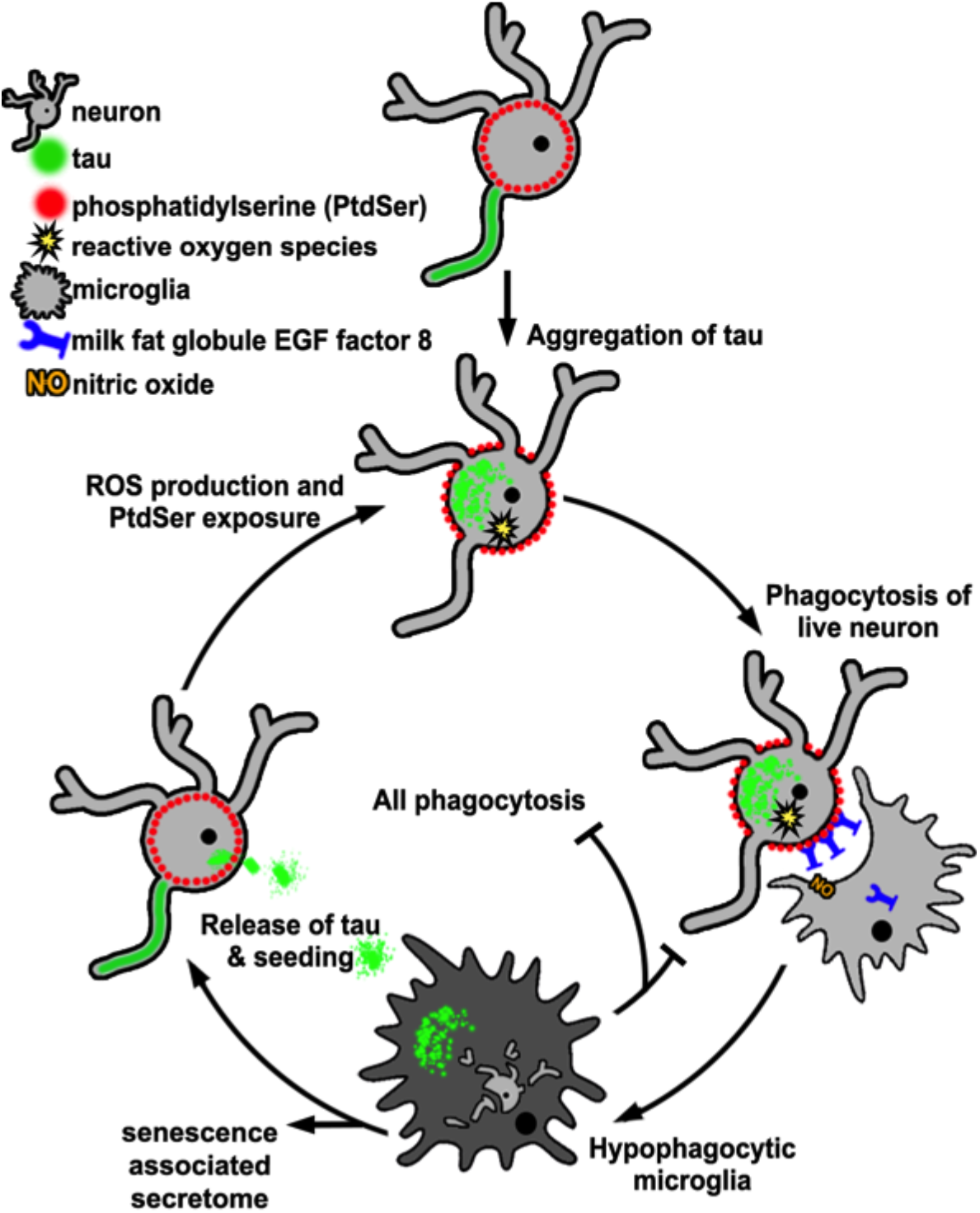
Schematic of proposed microglial dysfunction induced by phagocytosis of tau aggregate-containing neurons. Upon engulfment of neurons with tau aggregates, microglia become hypophagic, and acquire senescence-associated traits, resulting in accumulation of neurons with tau aggregates in a selfamplifying cycle.

Our data show that microglia that have phagocytosed neurons with insoluble tau aggregates not only maintain these aggregates for several days, but also expel aggregates capable of seeding tau aggregation. By contrast, monocultures of P301S DRGn containing tau aggregates did not expel tau, so tau release is unlikely to be a result of neuronal cell death or other factors in the culture environment. Our data showing an extended period of tau release are in agreement with a study where microglia isolated from adult rTg4510 mouse brains or AD brains also released tau over several days that was capable of seeding tau aggregation in a FRET-based biosensing cell assay (*47*). Interestingly, inflammatory stimuli failed to increase release of tau seeds from microglia in their study, so perhaps it is microglia with senescent phenotypes similar to those that we demonstrate here that are those most prone to release tau seeds. Microglia have previously been implicated as vectors of tau spreading *in vivo* through exosomes in mouse models with focal viral delivery of tau-GFP (*26*). However, no filamentous or misfolded tau was detected in these experiments, so it is not clear whether tau would be also be released by microglia via exosomes if it were highly aggregated, as is the case here. Although tau aggregate pathology is thought to develop along synaptically-connected routes through neuronally-released tau seeds (*24, 48*), there is evidence for lateral spread of tau as well (*49*). Thus it is possible that microglia surrounding neurons with tau pathology could act as sources of tau seeds, especially after PtdSer-dependent phagocytosis of tau aggregate-containing neurons or synapses that display PtdSer (*12, 13*). It is interesting that the presence of filamentous aggregated tau inside microglia in human tauopathy cases is a rare event (*50*), despite well-described cases of tau pathology in astrocytes and oligodendrocytes (*51, 52*). It is possible that microglia have a more efficient means of degrading or expelling tau aggregates (*53*) than astrocytes or oligodendrocytes. Another possibility is that microglia with tau aggregates fragment and die due to acquisition of a senescent phenotype (*37*) thereby releasing tau seeds. However, there was no noticeable microglial death in our system or in that of Hopp et al. (*47*) so the mechanism of tau aggregate release remains to be identified.

The present data demonstrate that microglia show hormetic behaviour: having been activated to engulf neurons with tau aggregates, they become hypophagocytic and show signs of senescence. The precise mechanism for this switch remains to be resolved. We tested whether PtdSer could be the signal since phagocytosis of live neurons by microglia depends on PtdSer exposure on the neuronal membrane (*4–6, 17, 54*), but hormesis was not mimicked by exposure of microglia to PtdSer liposomes, nor was MMP3 expression reduced by masking PtdSer with AnnV. The phenomenon of microglial hormesis after phagocytosis of β-amyloid has been described previously. For example, microglia that have phagocytosed β-amyloid fibrils showed reduced phagocytosis towards β-amyloid oligomers (*55*). Hypophagocytic behaviour was also observed *in vivo* in two mouse models with β-amyloid plaques (*56*) and demonstrated by the failure of freshly isolated microglia from 5xFAD transgenic mice to clear β-amyloid plaques from organotypic 5xFAD slice cultures whereas wildtype microglia displayed ample clearance (*57*). The switch from a pro-resolution phenotype to a hypophagocytic phenotype was suggested to represent a proinflammatory state that occurs when the microglia fail to digest and degrade engulfed highly stable filamentous protein aggregates (*58–60*). Nevertheless, exposure of microglia to extracellular monomeric or small oligomeric 2N4R tau increases phagocytosis (*4*) although this result is conditionally dependent on tau isoform, concentration, and phosphorylation status (*61*). The molecular differences between the different states of the microglia under these different settings remain to be resolved.

In addition to microglia being hypophagocytic, we show that coculture of microglia with P301S tau aggregate-containing neurons induced the hypofunctional microglia to secrete several senescence-associated factors (CXCL1, CXCL2, CCL5, IGFBP3, MMP9, and MMP3 (*62*)). The presence of dystrophic microglia that have been linked to senescence have been reported in Alzheimer’s disease and other neurodegenerative disorders (*63–66*) although no molecular signatures of senescence were recorded. Loss of phagocytic capacity, increased SA-β-gal activity, and altered cytokine profiles such as those we describe here are common signs of senescence. In this hypofunctional state microglia may lose their neuroprotective role and possibility exacerbate neurodegeneration. The mechanism of transduction to senescence is unclear although both cytokine signalling and phagocytic stress in retinal epithelium have been suggested to induce senescence (*67*). It will be important to identify the biochemical mechanism of these changes and determine if modification of the process would be beneficial in disease.

The production of MMP3 was one of the most notable changes induced by coculture of microglia with 5M P301S neurons. A recent transcriptomic analysis (*68*) describes an antiinflammatory microglial subtype that is induced by inflammation but this may not resemble the response we observe as none of the proteins we have identified in the CM are present as altered transcripts aside from CXCL10 and CCL2, both of which are linked to an inflammasome IFN signalling module. A weak link to phagocytosis is the upregulation of ARF and CDC42 transcripts, which modify the dynamics of actin polymerisation, but with these being more active, one may expect more pronounced phagocytosis rather than an inhibition such as the one we observe. Nevertheless, activation of MMP3 is a hallmark of several tauopathies, including AD and familial FTD cases (*69–71*). Since NFκB has been implicated in control of MMP3 expression in other cell types (*43, 72*), it was of interest to examine to what extent NFκB-dependent MMP3 production is linked to tau release from cocultured microglia. We found that the NFkB inhibitor BAY11-7082 prevented MMP3 production despite being applied under culture conditions that allow phagocytosis of the neurons, and therefore release of tau by microglia. MMP3 was still highly produced in the presence of AnnV, however, which blocks phagocytosis. Thus, NFkB appears to be activated as a result of cell signalling either between neurons with tau aggregates and microglia or between post-phagocytosis microglia and surrounding microglia. Since TNFa was not induced by the neurons, another possible mediator of NFkB activation are the reactive oxygen species produced by neurons with tau aggregates (*31, 73, 74*). The consequence of such strong up regulation of a matrix metalloprotease remains to be resolved. Loss of MMP3 represses the upregulation of other chemokines like CCL2, and CXCL1 (*75*), and could have effects at the synapse due to its tissue remodelling properties via the extracellular matrix (*76, 77*).

The key message from our study is that microglia switch to an alternative functional state as a result of coculture and phagocytosis of neurons with tau aggregates that is not classically inflammatory. Microglia that have phagocytosed tau aggregate-containing neuron produce tau seeds which may enhance tau pathology not only by spreading tau aggregation to new neurons but also by losing their normal phagocytic functions.

## Materials and Methods

### Experimental Design

The objective of these experiments was to test the functional state of microglia after phagocytosis of neurons containing insoluble filamentous hyperphosphorylated tau aggregates, and to determine the fate of the ingested tau. The study was designed as an *in vitro* cell culture system to investigate cellular and biochemical mechanisms leading to phenotypic changes in microglia and consequences for pathology, confirming key findings using brains from transgenic human P301S tau mice and human brain tissue from neurodegenerative diseases. We have previously published that dorsal root ganglion neurons from P301S mice with tau aggregates expose PtdSer and are phagocytosed live by cocultured microglia (*17*).

### Mice

Homozygous mice transgenic for human 0N4R P301S tau (henceforth P301S), and C57BL/6S (C57BL/6OlaHsd; henceforth C57) control mice were maintained as described previously (*29*). This research project was performed under the Animals (Scientific Procedures) Act 1986, Amendment Regulations 2012 following ethical review by the University of Cambridge Animal Welfare and Ethical Review Body (AWERB). Both female and male mice were used for experiments. Mice were housed in groups in individually ventilated cages, adding a plastic roll and nesting material for enrichment. Mice were kept under a 12 h light/dark cycle, with food and water available ad libitum.

### Human tissue

Human tissue was obtained from the Cambridge Brain Bank and the Alzheimer’s Centre at Indiana University. Handling of human tissue was according to the UK Human Tissue Act 2006 and is covered by the Cambridge Local Research Ethics Committee (LREC), approval number 09/40. Each case was neuropathologically confirmed as being either control, Pick’s disease, or expressing mutant P301L or +3 as described previously (*17, 80, 81*). The cohort contained mixed sexes and the age at death ranged from 53 to 76 (full details in Supplementary Table 1). Fresh frozen frontal cortex, or midbrain was homogenised in RIPA containing 2.5% SDS with protease inhibitors (Roche) at a ratio of 1:2 (w/v) with an IKA T-10 ULTRA TURRAX. Whole homogenates were clarified by centrifugation at 20,000 *xg* for 30 minutes at 4°C. The protein content in the resulting supernatant was determined by BCA assay (Pierce), and 5 μg protein was analysed by immunoblot.

### Cell cultures

DRG neurons (DRGn) from either P301S or C57 mice were cultured on 6-well culture plates coated with poly-D-lysine and laminin and maintained as described previously (*29*). Primary microglial cells were prepared from post-natal day 2-4 pups as previously described and cultured in DMEM containing 1% PSF (ThermoFisher) 10% heat inactivated FBS (ThermoFisher) and 50 ng/ml mouse colony stimulating factor 1 (mCSF1) (Peprotech). Neurons were cultured for 7 days in growth medium containing 20 μM fluorodeoxyuridine (to remove non-neuronal cells), then washed to remove residual debris and labelled in medium containing 3 μM pFTAA (a gift from K. Peter R. Nilsson, Linköping University, Sweden; (*31*)) for 30 min at room temperature to visualise tau aggregates. After washing the DRGn in PBS and replacing with coculture media, 200,000 primary microglia were added directly to the medium to create contact cocultures. Cells were cocultured in DMEM, 1% PSF, 5% heat inactivated FBS, 2 mM GlutaMax (Gibco) for 4 days at which time the conditioned medium (CM) was collected. Microglia were then re-isolated by physical dissociation in ice cold PBS without Mg2+ or Ca2+ (Gibco) and passed through a 40 μm pore membrane to exclude any contaminating DRGn, then plated onto uncoated 13 mm glass coverslips. Re-isolated microglia were either fixed in 4% PFA in PBS, or cultured in DMEM, 10% heat inactivated FBS, 1% PSF 10 ng/ml mCSF1 for a further 4 days. NFκB inhibition in microglia was performed by adding 5 μM BAY11-7082 (Sigma-Aldrich, B5556) for 1 h before microglial addition to DRGn. Annexin V (AnnV) (100 nM ImmunoTools 31490010) was added to the DRGn to pre-treat and mask PtdSer thus preventing phagocytosis.

### Seeding assay

HEK-P301S-venus cells were maintained in DMEM, 10% FBS and 1% PSF on uncoated glass 13mm coverslips. To assay tau aggregate seeding efficiency of CM, maintenance medium was replaced with conditioned medium for 48 h in the absence or presence of 1:1000 μl Lipofectamine 2000 (*33*). Cells were then fixed in 4% PFA and immunostained as below.

### Immunocytochemistry

Tau aggregates in P301S DRGn were labelled live with pFTAA as described previously (*31, 35*). Re-isolated microglia were counter stained with DAPI and or phalloidin-Alexa647 (LifeTechnologies), then mounted in Fluosave (MilliPore). HEK-P301S-venus cells were permeabilised in 0.3% Triton-X in PBS (PBST) and incubated with primary antibody overnight in PBST. Coverslips were washed in PBS and incubated with appropriate AlexaFluor-conjugated secondary antibody for 1 h at room temperature in PBST, counterstained with DAPI and mounted in FluorSave™. Images were taken on a Leica DMI 4000B microscope using a Leica DFC3000 G camera and the Leica application suite 4.0.0.11706. Images were analysed using ImageJ (Rasband, W.S., ImageJ, U.S. National Institutes of Health, Bethesda, Maryland, USA, http://imagej.nih.gov/ij/,1997-2014).

### ELISA

DRGn from P301S or C57 mice were cocultured with C57 microglia for 4 days to generate CM where tau was assayed. CM was first centrifuged at 20,000*g* for 20 mins to remove any cellular debris and the supernatant collected. Tau protein concentration in CM was assayed with antihuman tau ELISA according to manufacturer’s instructions (abcam, ab210972). Lactate dehydrogenase assay was performed on the CM according to the manufacturer’s instructions (abcam, ab102526). To assay the effect of NFκB inhibition, DRGn from P301S or C57 were cocultured with C57 microglia that had been pre-treated with BAY11-7082 or left untreated for 24 h to generate CM for MMP3 assay. MMP3 ELISA (abcam, ab100731) was performed on CM 24 h after addition of microglia according to manufacturers’ instructions.

### Immunoblotting

Cells were lysed in 1% NP40, 137 mM NaCl, 2 mM EDTA, 20 mM Tris HCl pH 8 with protease inhibitor cocktail (Roche) and assayed for protein concentration by the BCA assay (Pierce). Equal protein amounts were added to BioRad LDS loading buffer with 10% β-mercaptoethanol and run on 4-12% gradient SDS-PAGE gels. Proteins were transferred onto 0.2 μm pore PVDF membranes (Merck). Nonspecific background was blocked in 5% w/v dry skimmed milk (Sigma) in PBS containing 0.1% Tween 20 and membranes were incubated with the primary antibody (abcam, ab53015) overnight at 4°C, followed by 1 h at room temperature in the appropriate HRP-labelled secondary antibody (GE Healthcare). Blots were developed with ECL Clarity (GE Healthcare).

### Membrane filtration assay

CM were centrifuged at 20,000x*g* for 20 min to remove cellular debris. Cellulose acetate membranes (0.2 μm) were soaked in 5% SDS in dH2O for 10 min at room temperature, then mounted on to a 96 well vacuum chamber. Samples were incubated in 5% SDS in dH2O for 10 min at room temperature and 2 ml of each conditioned medium was filtered through the membrane under vacuum. Membranes were washed in 5% SDS under vacuum, rinsed in PBS and 0.1% tween20, then incubated with 2 μg/ml primary antibody (HT7) overnight at 4°C, and developed using the method described for immunoblotting. A sarkosyl-insoluble extract from 5 month-old P301S mouse spinal cord prepared as described previously (*30*) was the positive control. After initial signal development, each membrane dot was cut out of the membrane and bisected, half was treated with 70% formic acid (FA) for 15 min then dialysed over-night with Slide-a-lyzer (ThermoFisher) in PBS 0.1% tween20. The other half was minced, vortexed, and soaked in 5% SDS to elute available captured protein. Processed samples were then membrane filtered, antibody incubated and developed as before.

### Latex Bead phagocytosis assay

To assay phagocytic activity, fluorescent latex carboxylate-modified polystyrene beads, 2.0 μm mean particle size, were added to re-isolated microglial cultures on 13 mm coverslips at 1;10,000 dilution for 12 h at 37°C, or added to cultures of matched biological replicates pre-treated with either: 100 ng/ml LPS for 1 hour, 12 nM monomeric tau, 12 nM heparin-fibrillised tau (monomer equivalent), or 1 μg/ml PtdSer vesicles prepared by dissolving PtdSer in chloroform and drying under nitrogen before hydration in PBS and sonication to produce a vesicle suspension. Cultures were washed in PBS to remove free beads and fixed in 4% PFA. Microglia were counter stained with DAPI and phalloidin-Alexa647 and mounted with FluorSave™. Images were collected using the Leica DMI 4000B microscope and analyzed using ImageJ.

### SA-β gal assay

Re-isolated microglia were plated onto 13 mm coverslips and fixed in 4% PFA. The activity of β galactosidase at pH 6.0 was assayed as previously described (*78*). Briefly, a staining solution of 1 mg/mL 5-bromo-4-chloro-3-indolyl-beta-d-galactopyranoside (X-gal, Invitrogen), 1× citrate-sodium phosphate buffer (pH 6.0), 5 mM potassium ferricyanide, 5 mM potassium ferrocyanide, 150 mM NaCl, and 2 mM MgCl2 was applied to the fixed culture for 12 h at 37°C. Cultures were washed, counter stained with DAPI and mounted in FluorSave™. Mixed colorimetric and fluorescent images were collected using the Leica DMI 4000B microscope and analysed using ImageJ. Bleomycin was added as a positive control inducer of chemical senescence at 50 μg/ml for 12 hours.

### Cytokine array

CM was collected and centrifuged at 20,000 x*g* to remove cellular debris. Comparative cytokine amounts were assayed with the proteome profiler XL mouse cytokine array as per manufacturer’s instructions (R&D, ARY028). Densitometric measurements were quantified using ImageJ, and normalized between membranes using the membranes’ positive controls. To identify expression pathways, the data were analysed by STRING analysis ((*40*), https://string-db.org/). Interactions permitted were experimentally determined, co-expressed, and those that were predicted to be expressed by databases; any non-interacting proteins were removed from the analysis. Most confident partner interactions in the remaining array (threshold ≥ 0.9), yielding the following unique proteins: CXCL9, CXCL10, CXCL5, CXCL2, CXCL1, CCL5, CCL20, ENDOSTATIN, VEGFA, MMP2, CXCL13, IGFBP3, MMP9, PTX3, and MMP3. Noninteracting proteins were removed from analysis, as were Cxcl5, Cxcl9, Cxcl13, and Ccl20 due to lack of detectable expression across all conditions. The expression of 11 proteins across 3 biological replicates was expressed as a mean, standardised by calculating Z-scores, and then displayed on a heat-map.

### Principal Component analysis

PCA analysis was conducted using the Scikit-Learn library in Python (Version 3.7.6) (*79*). To calculate error bars (standard deviation), full replicate data (n=3) were used to ‘train’ the PCA transformation matrix, and error was calculated for each condition from the PC1 and PC2 coordinates of the replicates. This same transformation matrix was then applied to the mean-only dataset, and the error bars calculated previously were interpolated onto each corresponding condition.

### Statistical Analysis

Statistical analysis and graphing was performed using graph pad Prism version 7. All other samples were compared 1-way or 2-way analysis of variance (ANOVA) followed by an appropriate post hoc test, or by Students t test as indicated. Significant differences are reported as *p< 0.05, **p< 0.01, ***p< 0.001, ****p< 0.0001. Between 100-200 neurons were counted in each individual culture for determining the ratio of HT7/βIII tubulin.

## General

The authors would like to thank Michel Goedert (MRC Laboratory of Molecular Biology, Cambridge, UK) for his comments on the manuscript and use of the P301S transgenic mice. Human brain tissue was obtained from the Brain Library of the Dementia Laboratory, Department of Pathology and Laboratory Medicine at Indiana University School of Medicine and the Cambridge Brain Bank.

## Funding

This work was supported by an Alzheimer’s Research UK fellowship ARUK-RF2017A-4 to J.H.B., and grants from the European Union (EU/EFPIA/Innovative Medicines Initiative 2, Joint Undertaking n116060, IMPRIND (W.A.M, T.K., A.M.T, M.G.S), the Wellcome Trust and a Royal Society Sir Henry Dale Fellowship Grant Number 206248/Z/17/Z, (W.A.M), the UKDRI (funded by the UK Medical Research Council, Alzheimer’s Society and Alzheimer’s Research UK) (W.A.M, M.G.S), a iCase BBSRC and Eli Lilly studentship #G103374 (M.M., M.G.S). B.G. is supported by Department of Pathology and Laboratory Medicine, Indiana University School of Medicine and NIH grant P30-AG010133. The Cambridge Brain Bank is supported by the National Institute for Health Research (NIHR) Cambridge Biomedical Research Centre.

## Author contributions

JHB, AMT, and MGS conceived and designed research studies, JB performed the experiments, AMT assisted in the performance of the experiments, MM analysed the cytokine array data, TK provided the recombinant tau and assisted in experimental design, WAM provided the P301S tau-venus HEK cell line, MGS supervised project design, JB, MM, AMT and MGS interpreted the data and wrote the manuscript. All authors critically reviewed the manuscript.

## Competing interests

The authors have no competing interests to declare.

## Data and materials availability

All data needed to evaluate the conclusions in the paper are present in the paper and/or the Supplementary Materials

## Supplementary Materials

**Fig. S1.**
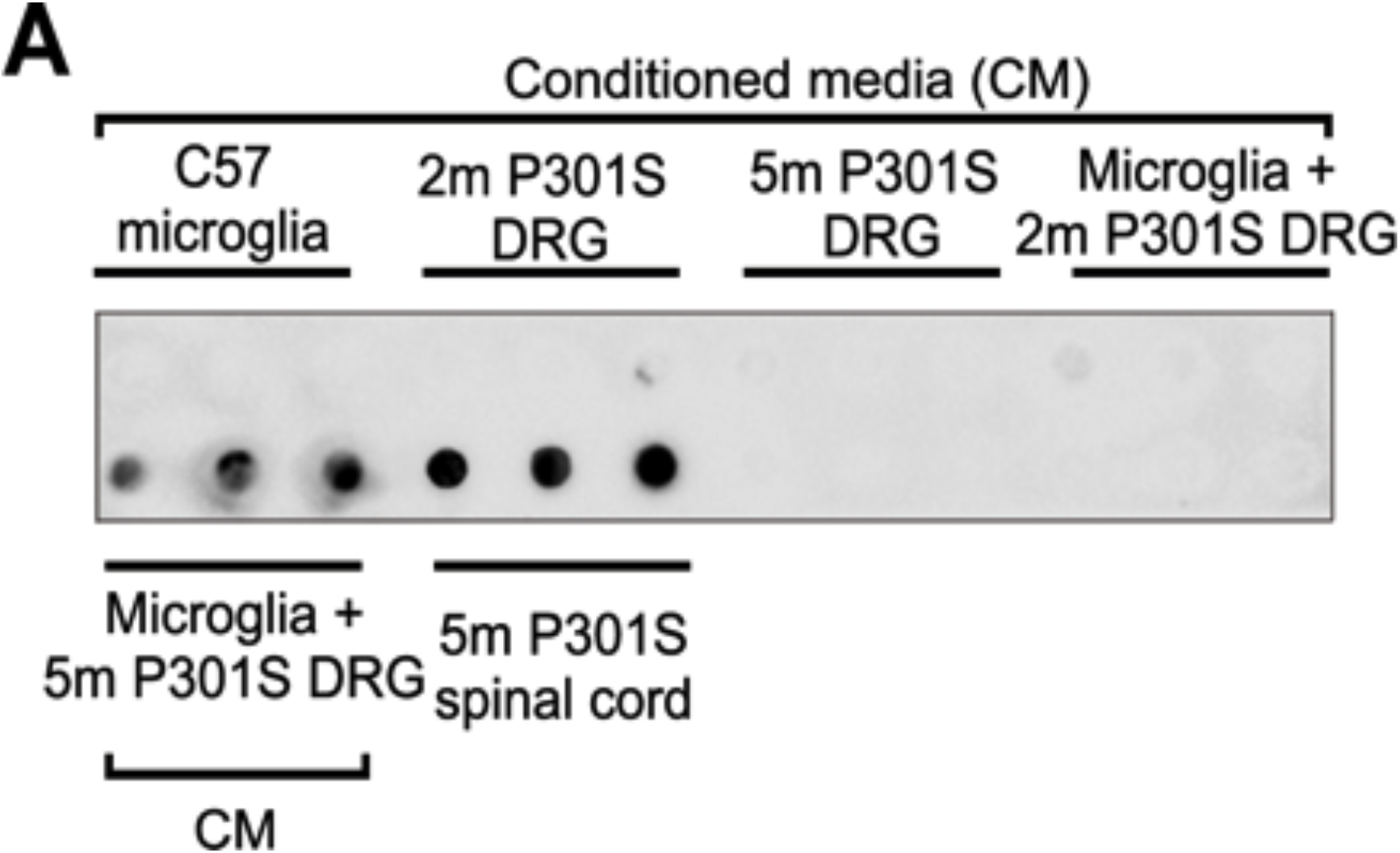
Intact blot with reference to Fig. 1. Original configuration of filter trap membrane in Figure 1D before segmenting, showing that all samples prior to extraction were processed and developed at the same time.

**Fig. S2.**
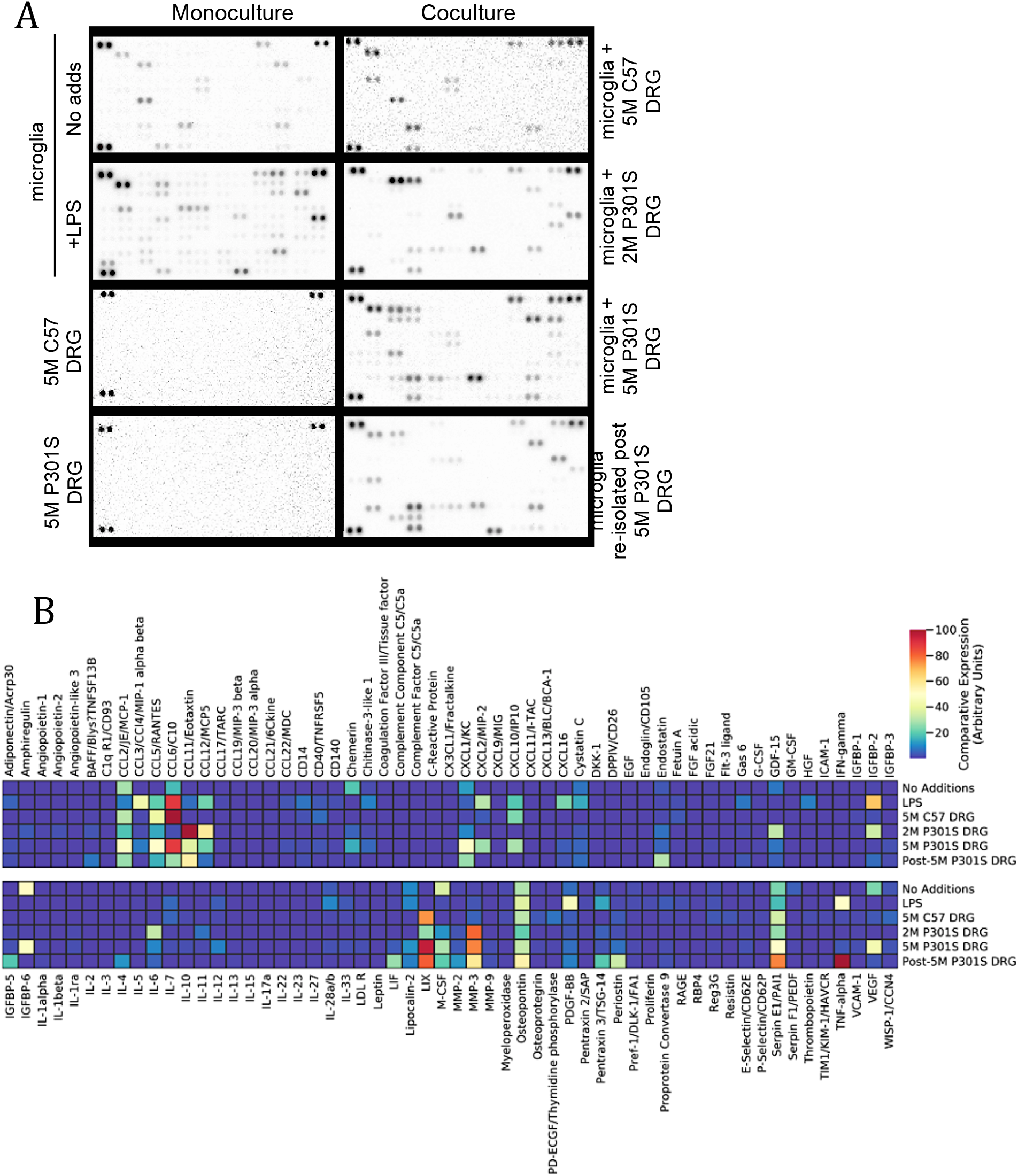
**A**) Examples of one set of membrane arrays probed with CM from (left column) monocultures of untreated microglia, microglia treated with LPS, pure DRG neurons from 5M C57 or P301S mice, and (right column) cocultures of microglia with DRG neurons from 5M C57, 2M P301S, or 5M P301S mice, and microglia isolated and recultured from cocultures with 5M P301S mice, used for quantification with reference to Fig. 3. The pair of spots on the top left, top right, and bottom left of each membrane are internal positive controls, and the right-most pair of (invisible) spots on the bottom right are internal negative controls. **B**) Relative expression heat map of total proteins on the cytokine arrays. The internal positive and negative controls are not represented.

**Supplementary Table 1.**
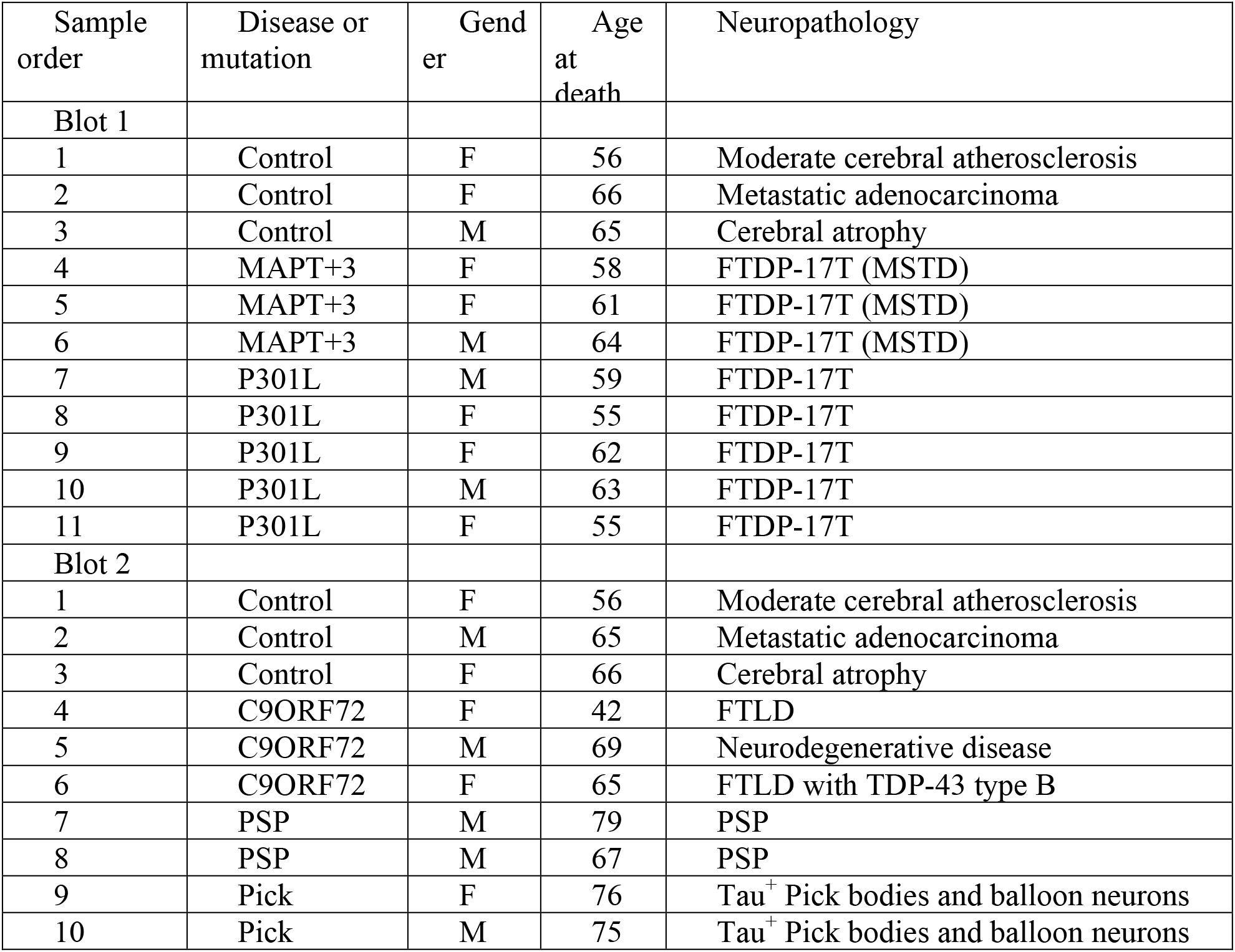
Human tissue details in relation to Fig. 4. Human brain tissue was obtained from The Alzheimer Disease Center, Indiana University School of Medicine and the Cambridge Brain Bank. Handling of human tissue was according to the UK Human Tissue Act 2006 and is covered by the Cambridge Local Research Ethics Committee (LREC), approval number 09/40. Grey matter (0.2 g) was mechanically homogenized in RIPA buffer containing 2.5% SDS with phosphatase and protease inhibitors at a 1:2 (w/v) ratio. Lysate was clarified at 20,000 g for 30 min and protein assayed. Abbreviations: FTDP-17T, frontotemporal dementia and Parkinsonism linked to chromosome 17, MSTD, Multiple system tauopathy with presenile dementia, FTLD, frontotemporal lobar degeneration. Controls are defined age matched subjects who did not present neurological disorders.

| Sample order | Disease or mutation | Gender | Age at death | Neuropathology |
| --- | --- | --- | --- | --- |
| Blot 1 |  |  |  |  |
| 1 | Control | F | 56 | Moderate cerebral atherosclerosis |
| 2 | Control | F | 66 | Metastatic adenocarcinoma |
| 3 | Control | M | 65 | Cerebral atrophy |
| 4 | MAPT+3 | F | 58 | FTDP-17T (MSTD) |
| 5 | MAPT+3 | F | 61 | FTDP-17T (MSTD) |
| 6 | MAPT+3 | M | 64 | FTDP-17T (MSTD) |
| 7 | P301L | M | 59 | FTDP-17T |
| 8 | P301L | F | 55 | FTDP-17T |
| 9 | P301L | F | 62 | FTDP-17T |
| 10 | P301L | M | 63 | FTDP-17T |
| 11 | P301L | F | 55 | FTDP-17T |
| Blot 2 |  |  |  |  |
| 1 | Control | F | 56 | Moderate cerebral atherosclerosis |
| 2 | Control | M | 65 | Metastatic adenocarcinoma |
| 3 | Control | F | 66 | Cerebral atrophy |
| 4 | C9ORF72 | F | 42 | FTLD |
| 5 | C9ORF72 | M | 69 | Neurodegenerative disease |
| 6 | C9ORF72 | F | 65 | FTLD with TDP-43 type B |
| 7 | PSP | M | 79 | PSP |
| 8 | PSP | M | 67 | PSP |
| 9 | Pick | F | 76 | Tau<sup>+</sup> Pick bodies and balloon neurons |
| 10 | Pick | M | 75 | Tau<sup>+</sup> Pick bodies and balloon neurons |

## Notes

### Competing Interest Statement

The authors have declared no competing interest.

